# Posterior white matter hyperintensities are associated with reduced medial temporal lobe subregional integrity and long-term memory in older adults

**DOI:** 10.1101/2022.09.19.508591

**Authors:** Batool Rizvi, Mithra Sathishkumar, Soyun Kim, Freddie Márquez, Steven J. Granger, Myra S. Larson, Blake A. Miranda, Martina K. Hollearn, Liv McMillan, Bin Nan, Nicholas J. Tustison, Patrick J. Lao, Adam M. Brickman, Dana Greenia, Maria M. Corrada, Claudia H. Kawas, Michael A. Yassa

## Abstract

White matter hyperintensities are a marker of small vessel cerebrovascular disease that are strongly related to cognition in older adults. Similarly, medial temporal lobe atrophy is well-documented in aging and Alzheimer’s disease and is associated with memory decline. Here, we assessed the relationship between lobar white matter hyperintensities, medial temporal lobe subregional volumes, and hippocampal memory in older adults.

We collected MRI scans in a sample of 139 older adults without dementia (88 females, mean age (SD) = 76.95 (10.61)). Participants were administered the Rey Auditory Verbal Learning Test (RAVLT). Regression analyses tested for associations among medial temporal lobe subregional volumes, regional white matter hyperintensities and memory, while adjusting for age, sex, and education and correcting for multiple comparisons.

Increased occipital white matter hyperintensities were related to worse RAVLT delayed recall performance, and to reduced CA1, dentate gyrus, perirhinal cortex (Brodmann area 36), and parahippocampal cortex volumes. These medial temporal lobe subregional volumes were related to delayed recall performance. The association of occipital white matter hyperintensities with delayed recall performance was fully mediated statistically only by perirhinal cortex volume.

These results suggest that white matter hyperintensities may be associated with memory decline through their impact on medial temporal lobe atrophy. These findings provide new insights into the role of vascular pathologies in memory loss in older adults and suggest that future studies should further examine the neural mechanisms of these relationships in longitudinal samples.

## INTRODUCTION

White matter hyperintensities (WMH) are regions of increased brightness that are best visualized on T2-weighted fluid-attenuated inversion recovery (FLAIR) magnetic resonance imaging (MRI) and are a radiological marker of small vessel cerebrovascular disease ^1^. WMH are associated with reduced cognitive function in older adults without dementia ^2,3^. Much of the work involving the contributions of WMH on cognitive functions in healthy older adults has thus far focused on the decline of processing speed and executive functioning ^3–5^, while the impact of WMH on memory specifically is less well characterized and studied ^6,7^. Importantly, WMH also contribute to both the onset and progression of Alzheimer’s disease (AD) and related pathophysiology ^8–11^. Supporting this line of work, WMH are implicated specifically in age- and AD-related neurodegeneration, providing some evidence of atrophy of medial temporal lobe (MTL) structures ^7,12–14^. However, the relationship between WMH and MTL subregional atrophy has been understudied.

The MTL plays a crucial role in episodic memory. In particular, the hippocampus and rhinal cortices are necessary for the encoding and consolidation of new episodic and semantic memories ^15,16^. Neurodegeneration of MTL subregions is linked to memory loss, both in aging and in AD. Hippocampal subfields including CA1 and the dentate gyrus (DG) undergo selective atrophy in aging and AD, changes that are, in turn, associated with substantial memory decline ^17–19^. Additionally, extra-hippocampal MTL cortical regions such as the perirhinal cortex and the entorhinal cortex are especially sensitive to effects of AD pathophysiology, including regional accumulation of tau pathology ^20–23^.

Despite evidence of both cerebrovascular related structural changes and MTL subregional atrophy in aging and AD, these two lines of work have been investigated mostly separately, and how these two features might be linked to memory decline has remained unclear. Additionally, while most of the work on WMH and cognition applies measures of global WMH burden, other studies point to the utility of investigating the effects of regionally specific WMH on cognition ^6,13,24,25^. Likewise, the study of MTL atrophy subregionally, including hippocampal subfields and extra-hippocampal subregions, allows us to further understand these regionally specific associations ^23,26^

In the current study, we first determined whether regional WMH accumulation is associated with memory outcomes. We focused on word list delayed recall performance as assessed by the Rey Auditory Verbal Learning Test (RAVLT). We tested the hypothesis that regional patterns of WMH accumulation will be associated with MTL subregional volumes in older adults without dementia, and subsequently tested whether these MTL subregional volumes are related to delayed recall performance. Finally, using a mediation model, we tested the hypothesis that the association of WMH with memory performance is through their impact on MTL subregional volume.

## MATERIALS AND METHODS

### Participants

One-hundred thirty-nine participants were included in the study. Eighty-seven community-dwelling adults were part of the Biomarker Exploration in Aging, Cognition, and Neurodegeneration (BEACoN) study, 23 were recruited from the UCI Alzheimer’s Disease Research Center (ADRC) longitudinal cohort, and 29 were recruited from the 90+ Study of the Oldest Old. All participants gave written informed consent in agreement with the Institutional Review Board of the University of California. The study only included participants who underwent both MR imaging (T2- and T1-weighted MRI) and neuropsychological testing.

Participants were included if they had performed RAVLT and had structural images analyzed. A total of N=139 participants with RAVLT measures had WMH (including n=130, missing n=9) and/or MTL volume data (included n=134, missing n=5). None of the participants were diagnosed with dementia at the time of testing. However, a subset of them (n=18) received a diagnosis of cognitive impairment (either mild cognitive impairment – MCI (n=4), cognitive impairment/no dementia – CIND (n=6), or questionable cognitive impairment – QCI (n=8)). This subset of participants is color coded in all association plots. Demographic characteristics of the participants are included in Table 1.

**Table 1.**
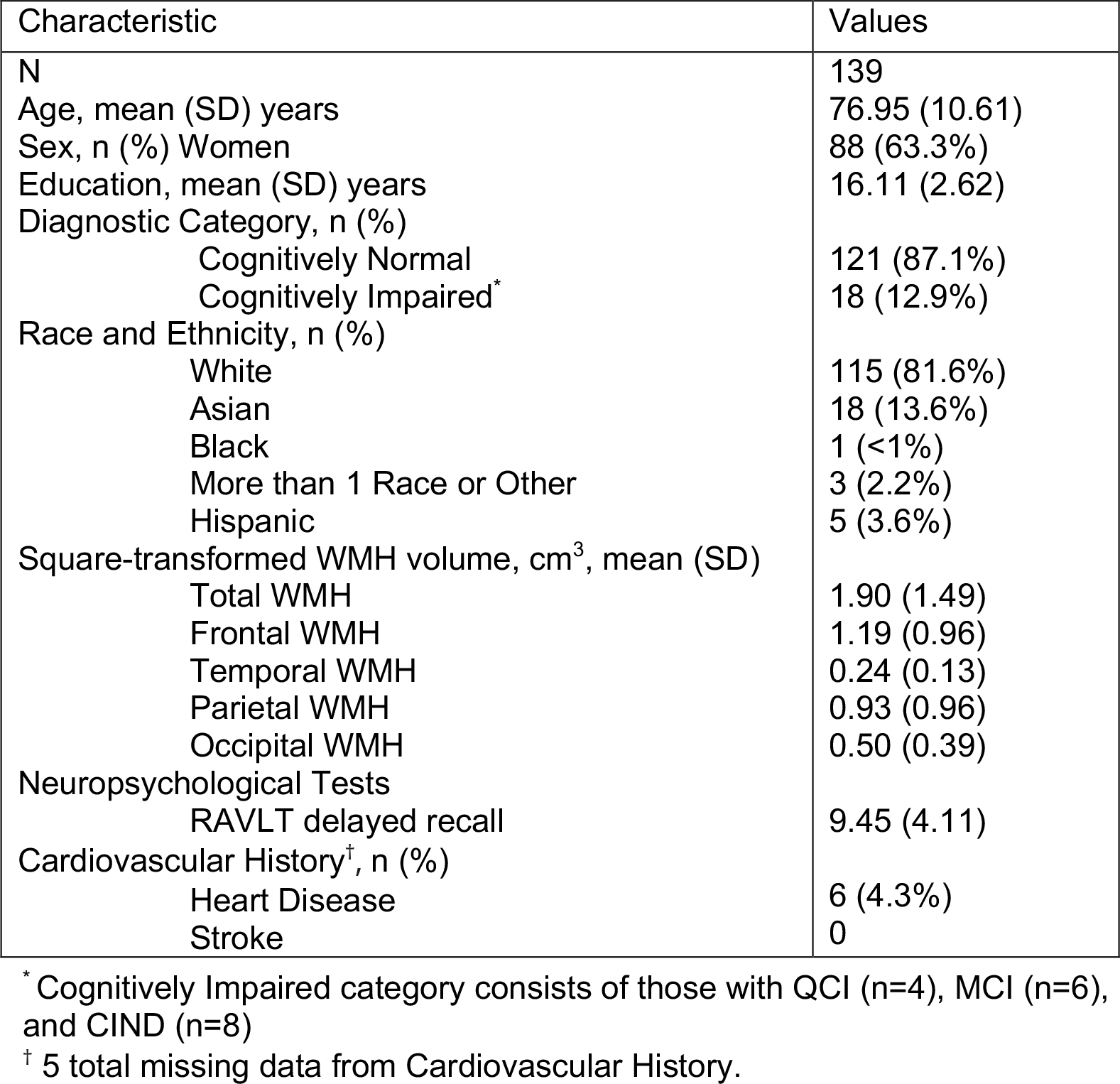
Demographics and Summary Variables

| Characteristic | Values |
| --- | --- |
| N | 139 |
| Age, mean (SD) years | 76.95 (10.61) |
| Sex, n (%) Women | 88 (63.3%) |
| Education, mean (SD) years | 16.11 (2.62) |
| Diagnostic Category, n (%) |  |
| Cognitively Normal | 121 (87.1%) |
| Cognitively Impaired* | 18 (12.9%) |
| Race and Ethnicity, n (%) |  |
| White | 115 (81.6%) |
| Asian | 18 (13.6%) |
| Black | 1 (<1%) |
| More than 1 Race or Other | 3 (2.2%) |
| Hispanic | 5 (3.6%) |
| Square-transformed WMH volume, cm<sup>3</sup>, mean (SD) |  |
| Total WMH | 1.90 (1.49) |
| Frontal WMH | 1.19 (0.96) |
| Temporal WMH | 0.24 (0.13) |
| Parietal WMH | 0.93 (0.96) |
| Occipital WMH | 0.50 (0.39) |
| Neuropsychological Tests |  |
| RAVLT delayed recall | 9.45 (4.11) |
| Cardiovascular History<sup>†</sup>, n (%) |  |
| Heart Disease | 6 (4.3%) |
| Stroke | 0 |

\* Cognitively Impaired category consists of those with QCI (n=4), MCI (n=6), and CIND (n=8)
<sup>†</sup> 5 total missing data from Cardiovascular History.

### Magnetic Resonance Imaging

For the BEACoN and 90+ cohorts, magnetic resonance imaging (MRI) data were acquired on a 3.0 Tesla Siemens Prisma scanner at the Facility for Imaging and Brain Research (FIBRE) at the University of California, Irvine. The following scans were acquired: Structural T1-weighted MPRAGE (resolution = 0.8 × 0.8 × 0.8 mm, repetition time = 2300 ms, echo time = 2.38 ms, FOV read = 256 mm, slices = 240, slice orientation = sagittal), T2-weighted fluid-attenuated inversion recovery (FLAIR; resolution = 1.0 × 1.0 × 1.2 mm, repetition time = 4800 ms, echo time = 441 ms, FOV read = 256 mm, slices = 160, inversion time = 1550 ms, slice orientation = sagittal) and T2-Turbo Spin Echo (resolution = 0.4 mm × 0.4 mm × 2.0, repetition time = 5000 ms, echo time = 84 ms, FOV read = 190 mm). For the ADRC cohort, MRIs were acquired on a 3 Tesla Philips scanner. The following scans were acquired: Structural T1-weighted MPRAGE (resolution = 0.54 × 0.54 × 0.65 mm, repetition time = 11 ms, echo time = 18 ms, FOV = 240 mm x 231 mm × 150 mm, slices = 231, slice orientation = sagittal), T2-weighted FLAIR (FLAIR; resolution = 0.65 × 0.87 × 4 mm, repetition time = 11000 ms, echo time = 125 ms, FOV = 230 mm × 103 mm × 119, slices = 24, inversion time = 2800 ms, slice orientation = transverse) and T2-Turbo Spin Echo (resolution = 0.47 mm × 0.47 mm × 2 mm, repetition time = 3000 ms, echo time = 80 ms, FOV = 180 mm × 180 mm × 109 mm).

### MRI analyses

#### White matter hyperintensities segmentation

Image processing leveraged the open-source ANTsX software ecosystem ^32^ with a particular focus on specific deep learning applications developed for neuroimaging made available for both Python and R via the ANTsXNet (ANTsPyNet/ANTsRNet) libraries. Specifically, for the work described here, WMH segmentation and lobar parcellation (see Figure 1) based on the Desikan-Killiany-Tourville (DKT) cortical labels ^33^ employed the two ANTsPyNet functions respectively: `sysu_white_matter_hypterintensity_segmentation` and `desikan_killiany_tourville_labeling`.

**Figure 1.**
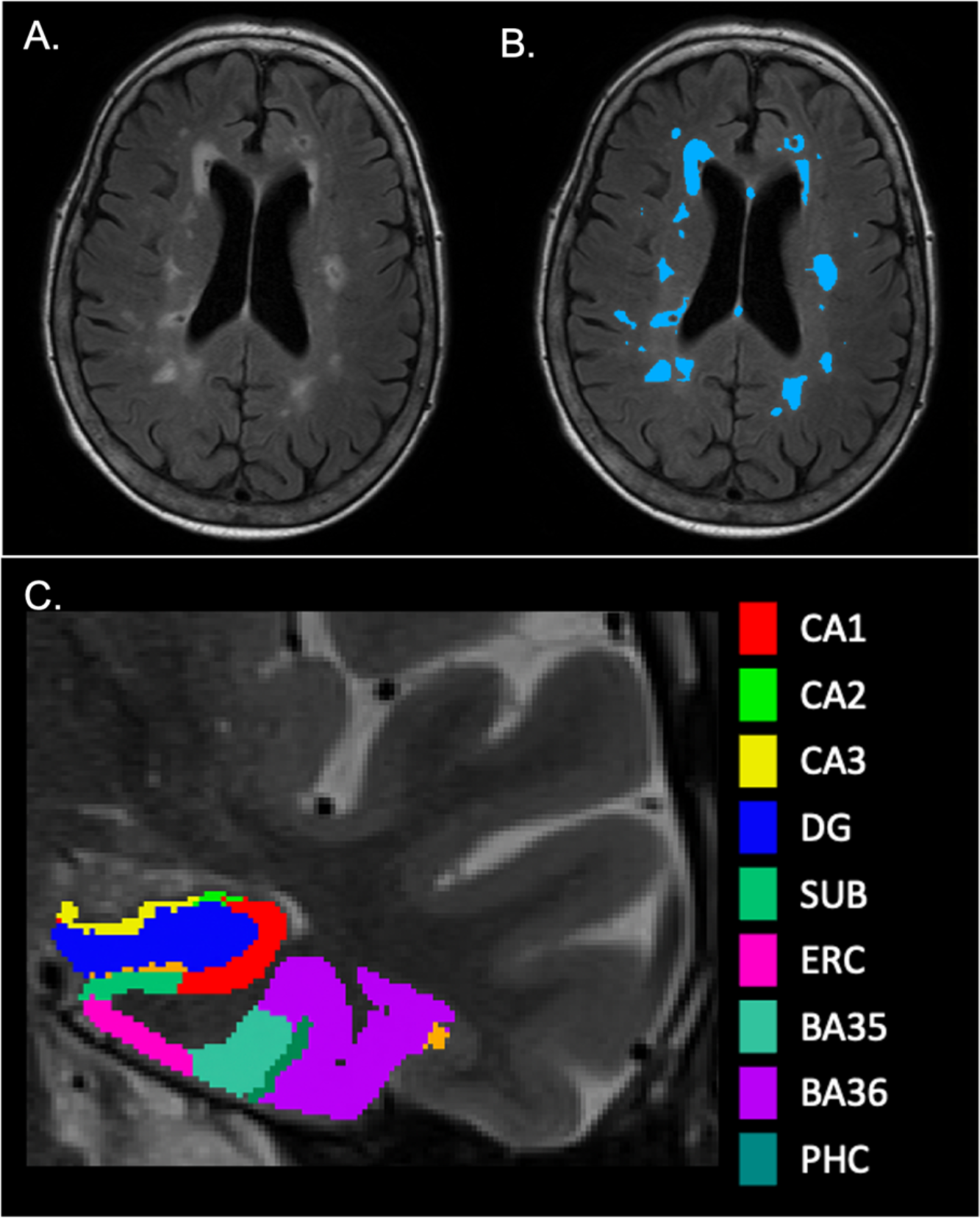
**(A)**. Axial view of a FLAIR image with unlabeled WMH. **(B)**. Axial view of a FLAIR image with labeled WMH. **(C)**. Coronal view of a T2-weighted image with MTL segmentation from ASHS. PHC is not captured in this image slice.

In conjunction with the International Conference on Medical Image Computing and Computer Assisted Intervention (MICCAI) held in 2017, a challenge was held for the automatic segmentation of WMH using T1-weighted and FLAIR images ^34^. The winning entry used a simplified preprocessing scheme (e.g., simple thresholding for brain extraction) and an ensemble (n=3) of randomly initialized 2-D U-nets to produce the probabilistic output ^35^. Importantly, they made both the architecture and weights available to the public. This permitted a direct porting to the ANTsXNet libraries with the only difference being the substitution of the threshold-based brain extraction with a deep-learning approach ^32^. All WMH masks resulting from the above automated segmentation procedure were manually edited by an expert rater (B.R.) for improved accuracy.

The segmentation above was followed by lobar parcellation. The process involved an automated, deep learning-based DKT labeling protocol for T1-weighted images, which was described in Tustison ^32^ in the context of cortical thickness. Briefly, data from several neuroimaging studies described in Tustison ^36^ were used to train two deep learning networks—one for the “inner” (e.g., subcortical, cerebellar) labels and one for the “outer” cortical labels. After an individual T1-weighted scan was labeled with the cortical DKT regions, the six-tissue (i.e., CSF, gray matter, white matter, deep gray matter, cerebellum, and brain stem) segmentation network was applied to the skull stripped image. Cortical labels corresponding to the same hemispheric lobes were combined and then propagated through the non-CSF brain tissue to produce left/right parcellations of the frontal, temporal, parietal, and occipital lobes, as well as left/right divisions of the brainstem and cerebellum. Left and right lobar WMH volumes were derived and summed across hemispheres. Due to a positively skewed distribution and values of zero within our WMH data, we square root transformed all regional and total WMH data.

#### MTL subregional volumes

Medial temporal lobe subregions including hippocampal subfields were automatically segmented with the Automatic Segmentation of Hippocampal Subfields (ASHS) ^37^ software using T1 and T2-weighted images. The ASHS pipeline implements joint label fusion and corrective learning to accurately segment hippocampal subfield volumes and cortical medial temporal lobe subregions. The resulting output included volumes of the following subregions in native T2 space: CA1, CA2, CA3, dentate gyrus (DG), subiculum, entorhinal cortex (ERC), perirhinal cortex subdivided into Brodmann Areas 35 and 36 (PRC; BA35, BA36), and parahippocampal cortex (PHC) volumes (see Figure 1). Left and right subregional volumes were summed to provide total subregional volumes. The resulting total subregional volumes were then adjusted for total intracranial volume (TIV), by dividing the total subregional volume by the individual’s TIV. The ratios were then multiplied by 1000: (MTL subregional volume / TIV) x 1000. TIV was obtained by implementing ANTs brain extraction and creating a binary brain mask and calculating total volume of the brain mask.

### Neuropsychological Testing

Participants were administered the Rey Auditory Verbal Learning Test (RAVLT), which assesses word list learning and memory, including rate of learning, retention, and recognition memory. RAVLT delayed recall has is sensitive to age-related memory decline ^38,39^, and thus was used as the outcome measure of primary interest. Other outcome measures of RAVLT, including learning slope, percent forgetting, retroactive interference and recognition, are reported in supplemental tables.

### Statistical Analysis

Statistical analyses were performed using SPSS Statistics v. 28. The first three sets of analyses below were performed using linear regressions. In the first set of analyses, we tested the separate associations between regional and total WMH and RAVLT delayed recall scores. We report associations between regional and total WMH and other RAVLT outcome measures in Supplemental Table 1. In the second set of analyses, we tested associations between regional WMH and total MTL subregional volumes in separate models. For associations found to be significant between regional WMH and MTL total MTL subregional volumes, we report associations with the regional WMH and lateralized (left and right) MTL subregional volumes in Supplemental Table 2. MTL subregions that were significantly associated with regional WMH were subsequently included in the third set of analyses to separately test the association between the specific MTL subregions and RAVLT delayed recall. We report associations with between specific MTL subregions and other RAVLT outcomes measures in Supplemental Table 3. The associations between the lateralized (left and right) MTL subregional volumes and RAVLT delayed recall are reported in Supplementary Table 4. All regression models adjusted for age, sex, and education. For analyses testing associations between regional WMH and MTL subregions, multiple comparisons were corrected for using the Holm-Bonferroni method. We then conducted mediation models that tested the effect of regional WMH on delayed recall, with MTL subregional volume(s) mediating this relationship, while adjusting for age, sex, and education. Regional WMH that were associated with MTL subregional volumes were included as the independent variable; MTL subregional volumes that were associated with delayed recall were included as the mediator(s). As post-hoc analyses, we included study cohort as an additional covariate in the above models.

## RESULTS

One-hundred thirty-nine participants (88 F, mean age = 76.95 ± 10.61) were included in the study. Table 1 displays their demographic and other summary characteristics.

### Associations between regional WMH and memory

We tested whether total and regional (lobar) WMH were associated with RAVLT delayed recall, while adjusting for age, sex, and education. We found that only increased occipital WMH were related to lower RAVLT delayed recall scores (*b* = −2.009, 95 CIs [−3.929, −0.089], p = 0.040; Figure 2A). When adding cohort as an additional covariate, the effect size remained similar (*b* = −1.730)

**Figure 2.**
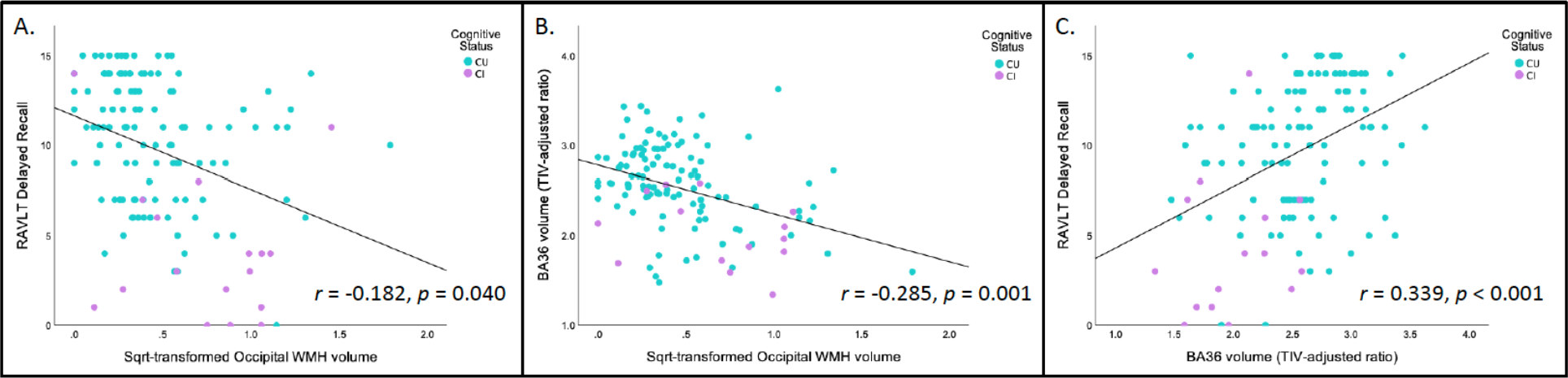
In all three scatterplots, cognitive status is color-coded, with cognitively unimpaired (CU) in blue and cognitively impaired (CI) in purple. **(A)** A scatterplot of the negative association between square root transformed occipital WMH and delayed recall. **(B)**. A scatterplot of the negative association between square root transformed occipital WMH and BA36 volume (TIV adjusted ratio). **(C)**. A scatterplot of the positive association between BA36 volume (TIV adjusted ratio) and delayed recall. Other significant associations found that were not involved in the mediation model (as seen in Figure 3) can be found in Supplemental materials

### Associations between regional WMH and MTL subregional volumes

We examined associations between total and regional WMH and MTL subregional volumes, while adjusting for age, sex, and education. After applying Holm-Bonferroni multiple comparison correction, increased occipital WMH was associated with reduced CA1 (*b* = −0.254, 95% CIs [−0.408, −0.109=0], p = 0.001), DG (*b* = −0.163, 95% CIs [−0.256, −0.069], p < 0.001), BA36 volumes (*b* = −0.426, 95% CIs [−0.686, −0.167], p = 0.001; Figure 2B), and PHC volumes (*b* = −0.212, 95% CIs [−0.366, −0.059], p = 0.007). No other regional or total WMH associations with MTL volume survived Holm-Bonferroni correction. Effect sizes of our findings remained similar once cohort was added as an additional covariate (CA1: *b* = −0.247, DG: *b* = −0.156, BA36: *b* = − 0.410, PHC: *b* = −0.202)

### Associations between MTL subregional volumes and delayed recall memory

MTL subregions that were significantly associated with WMH after applying Holm-Bonferroni correction were included in the second analysis, in which we tested associations between MTL subfield volumes and RAVLT delayed recall. All four MTL subregional volumes, including CA1, DG, BA36, and PHC, were also significantly associated with RAVLT delayed recall performance (CA1: *b* = 2.114, 95% CIs [0.115, 4.113], p = 0.038; DG: *b* = 3.956, 95% CIs [0.737, 7.176], p = 0.016; BA36: *b* = 2.445, 95% CIs [1.263, 3.628], p < 0.001; PHC: *b* = 2.381, 95% CIs [0.222, 4.541], p = 0.031; Figure 2C; Effect sizes of our findings remained similar once cohort was added as an additional covariate (CA1: *b* = 1.775; DG: *b* = 3.216; BA36: *b* = 2.201; PHC: *b* = 1.851)

### MTL subregional volumes mediate the effect of WMH on delayed recall memory

Four separate mediation models were run, informed by the previous analyses. As only occipital WMH were associated with memory and with MTL subregional volumes, it was included in the mediation model as the primary independent variable. CA1, DG, BA36, and PHC volumes were associated with both occipital WMH and with delayed recall, and thus were included as mediators separately in the four models. We found there was no direct effect of occipital WMH on delayed recall in any mediation model (M=CA1: b = −1.513, p = 0.150, 95% CIs [−3.580, 0.554]; M=DG: b =−1.399, p = 0.183, 95% CIs [−3.467, 0.668]; M=BA36: b = −0.895, p = 0.386, 95% CIs [−2.837, 1.106]; M=PHC: b = −1.528, p = 0.140, 95% CIs [−3.565, 0.506]). However, there was an indirect effect of occipital WMH on delayed recall and that only BA36 volume mediated this association (M=BA36: b = −1.052, 95% CIs [−2.184, −0.233]; Figure 3). The effect size of the indirect effect of occipital WMH on delayed recall with BA36 volume mediating the association was similar when cohort was included as an additional covariate (M=BA36: b = − 0.8905).

**Figure 3.**
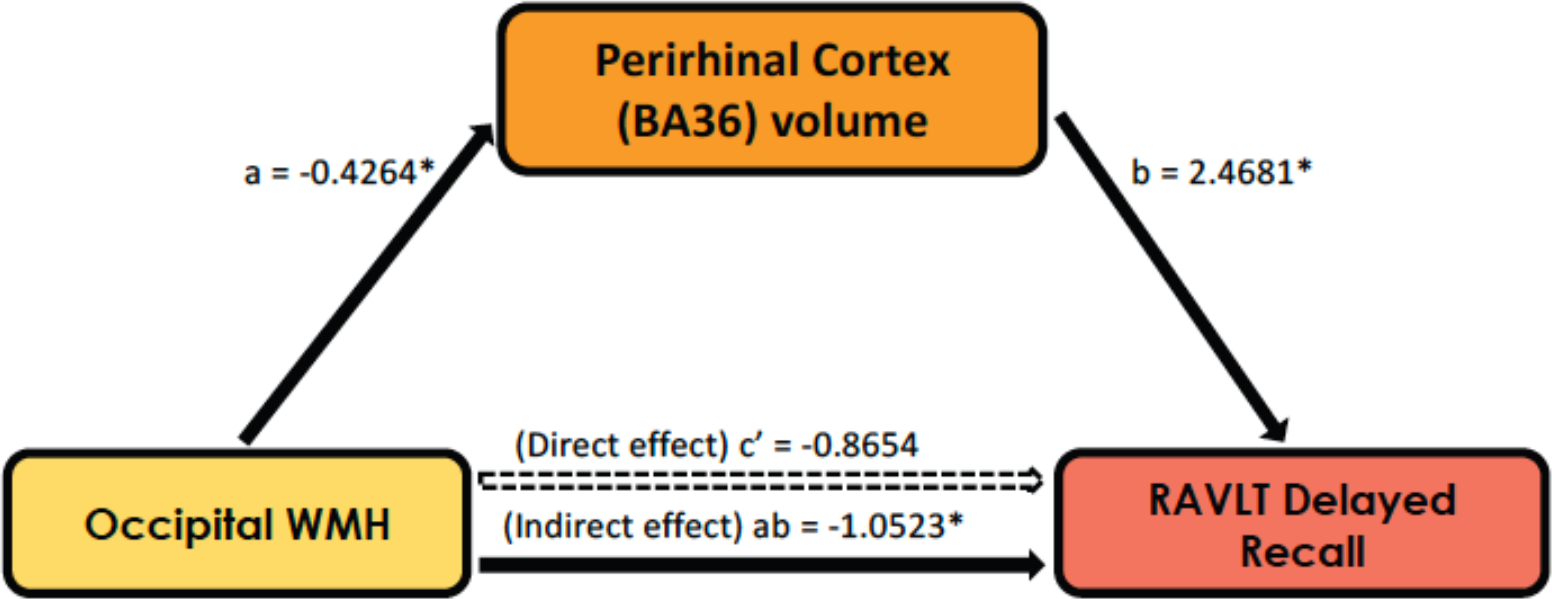
In this mediation model, there is an indirect effect where perirhinal cortex (BA36) volume mediates the relationship between occipital WMH and RAVLT delayed recall. Path a: *b*=−0.426, p=0.0015; Path b: *b*=2.4681, p=0.0003; Path c’ (Direct effect): *b*=−0.8654, 95% CIs [−2.8366, 1.1058]; Path ab (Indirect effect): *b*=−1.0523, 95% CIs [−2.1835, −0.2328].

## DISCUSSION

We interrogated the relationships among regional WMH, MTL subregional volumes, and delayed memory in older adults. We found that only occipital WMH burden was associated with lower RAVLT delayed recall performance. Increased occipital WMH was also associated with reduced CA1, DG, perirhinal cortical (BA36), and parahippocampal cortical (PHC) volumes. CA1, DG, perirhinal cortical (BA36) and parahippocampal volumes were related to worse RAVLT delayed recall in these older adults. Furthermore, in a mediation test, we found that occipital WMH had an indirect effect on delayed recall with perirhinal cortex (BA36) volume fully mediating this association.

Our results provide new evidence that regional patterns of small vessel cerebrovascular disease, here measured as lobar WMH volume, exhibits associations with medial temporal and hippocampal subfield volumes in older adults without dementia. Increased occipital WMH volume was related to lower volumes of CA1, DG, perirhinal cortex (BA36), and parahippocampal cortex (PHC) – all of which are structures that have been found to be critical to episodic memory formation and retrieval in older adults ^40,41^. Our results are somewhat in contrast to a recent study ^42^ suggesting that WMH were related to longitudinal atrophy of the subiculum in aging participants, although they did not relate this association to neuropsychological outcome measures. Another study in older adults with depression showed that depressed older individuals had smaller perirhinal (BA36) volumes and that higher temporal WMH volume was related to lower BA36 volume ^43^.

The DG is known to be important for pattern separation – the ability to store similar experiences using non-overlapping neural codes ^44^, which is known to be compromised with aging and AD. The CA1 is the hippocampus’ major output pathway and is also known to be impacted by aging and AD ^45^. Similarly, the parahippocampal and perirhinal cortices undergo early age-related structural changes and have important implications for episodic memory decline ^46,47^.

Importantly, the perirhinal cortex and the transentorhinal regions are among the first to deposit tau pathology in AD ^48^, and regional accumulation of tau in the temporal lobes as measured by tau PET is correlated with atrophy of the perirhinal cortex ^22^. Mechanistically, it is possible that WMH may increase susceptibility to neurodegeneration in areas typically affected in AD such as the MTL via their role in promoting tau hyperphosphorylation ^9^.

A potentially surprising aspect of our findings was that only occipital WMH of the regionally subdivided WMH volumes was related to lower MTL and hippocampal subfield volumes. While we may have not expected this pattern to be restricted to occipital WMH, previous work has indeed demonstrated that posterior WMH, including parietal and occipital WMH, are more linked to earlier AD onset in older adults ^49^. Additionally, posterior WMH has been associated with greater entorhinal cortical thinning, along with an increase in longitudinal CSF tau ^13,14^. The neurobiological mechanism by which posterior WMH contribute more to cognitive decline is yet to be elucidated. One possible mechanism could be related to the vascular supply to these regions. As the posterior cerebral artery (PCA) supplies blood to posterior areas, hippocampus, and areas of the MTL ^27^, a possible explanation may be that lower perfusion leads to increased posterior WMH, which has also been linked to tau accumulation ^9,14^ and subsequent MTL neurodegeneration ^50^.

There were some limitations of this study that are important to note. First, this study was conducted in a convenience sample that is comprised of predominantly affluent, highly educated, non-Hispanic white participants, which is not representative of an ethnically and socioeconomically diverse population. Second, due to a cross-sectional study design, this study cannot infer longitudinal relationships. Lastly, we did not have the ability to examine how clinical diagnosis would interact with these associations, due to a very limited number of individuals with mild cognitive impairment and the absence of patients with dementia. Future work should attempt to incorporate data from more diverse samples, increase the representation across the clinical impairment spectrum, as well as follow participants longitudinally to monitor the onset of these biomarker features over time. Additional inclusion of markers of AD neurodegenerative pathology such as tau PET will also help understand whether spatial colocalization of tau and MTL atrophy occurs as a result of posterior WMH.

In summary, the current work demonstrates the potential role of WMH in MTL atrophy, specifically increased posterior WMH burden in relation to lower volume of hippocampal subfields and surrounding cortex. The study specifically identified an association between occipital WMH and word list delayed recall, whereby perirhinal cortex (BA36) volume fully mediated this association. This work highlights the relevance of cerebrovascular pathology to neurodegenerative changes and cognitive decline even in older adults without dementia. It also suggests that examining modifiable vascular risk factors that can lower cerebrovascular burden could potentially reduce or slow down neurodegenerative cascades and thereby stem memory loss in older adults, particularly those at increased risk for AD.

## COMPLETING INTERESTS

M.A.Y. is co-founder and chief scientific officer of Augnition Labs, LLC.

