## Supplementary Material for "Posterior white matter hyperintensities are associated with reduced medial temporal lobe subregional integrity and long-term memory in older adults"

Rizvi et al.

### SUPPLEMENTARY FIGRUES

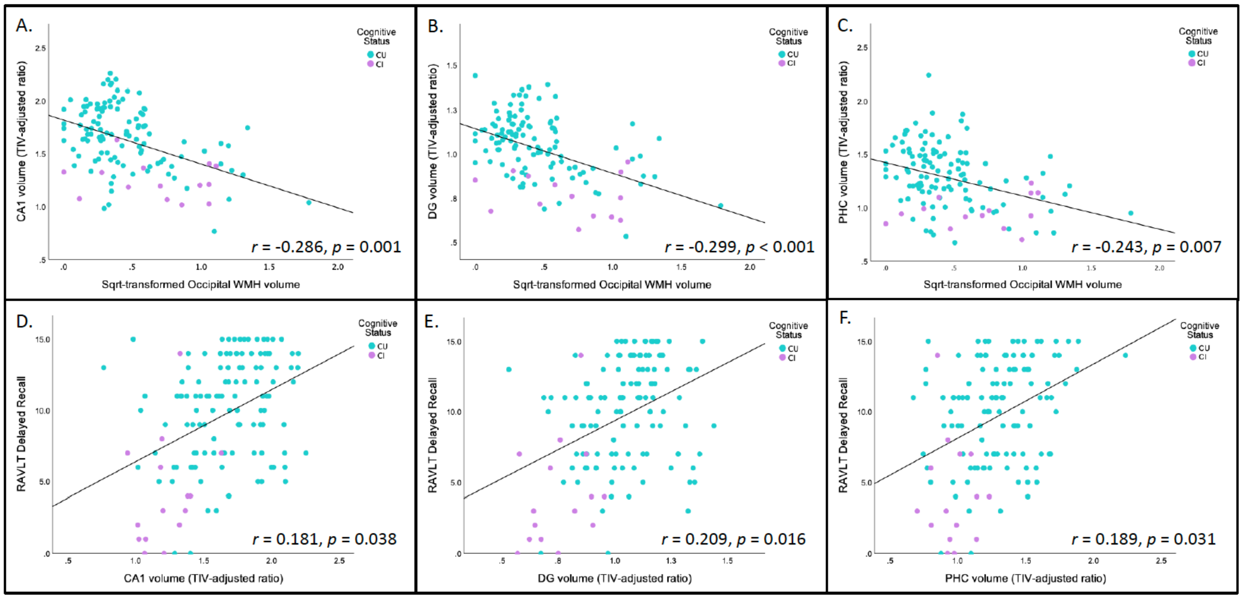


**Supplemental Figure 1. A.** A scatterplot of the negative association between square root transformed occipital WMH and CA1 volume (TIV adjusted ratio). **B.** A scatterplot of the negative association between square root transformed occipital WMH and DG volume (TIV adjusted ratio). **C.** A scatterplot of the negative association between square root transformed occipital WMH and PHC volume (TIV adjusted ratio). **D.** A scatterplot of the positive association between CA1 volume (TIV adjusted ratio) and delayed recall. **E.** A scatterplot of the positive association between DG volume (TIV adjusted ratio) and delayed recall. **F.** A scatterplot of the positive association between PHC volume (TIV adjusted ratio) and delayed recall.

### SUPPLEMENTARY TABLES

**Supplementary Table 1.** Associations between WMH and RAVLT measures

| WMH Distribution | RAVLT Measure | *b*-coefficient | Standardized β-coefficient | p-value | 95% CI |
|---|---|---|---|---|---|
| Frontal WMH | Delayed Recall | 0.273 | 0.069 | 0.407 | (-0.377, 0.924) |
| | Learning Slope | 0.087 | 0.149 | 0.116 | (-0.022, 0.195) |
| | Percent Forgetting | -0.016 | -0.065 | 0.475 | (-0.062, 0.029) |
| | Retroactive Interference | 0.003 | 0.013 | 0.897 | (-0.041, 0.047) |
| | Recognition | 0.054 | 0.035 | 0.355 | (-0.248, 0.356) |
| Temporal WMH | Delayed Recall | 0.605 | 0.044 | 0.574 | (-1.518, 2.728) |
| | Learning Slope | 0.127 | 0.063 | 0.484 | (-0.230, 0.483) |
| | Percent Forgetting | -0.011 | 0.013 | 0.883 | (-0.160, 0.137) |
| | Retroactive Interference | 0.008 | 0.010 | 0.912 | (-0.135, 0.151) |
| | Recognition | -0.087 | -0.016 | 0.862 | (-1.070. 0.897) |
| Parietal WMH | Delayed Recall | 0.104 | 0.024 | 0.777 | (-0.624, 0.832) |
| | Learning Slope | 0.070 | 0.111 | 0.255 | (-0.051, 0.192) |
| | Percent Forgetting | 0.004 | 0.013 | 0.890 | (-0.047, 0.054) |
| | Retroactive Interference | -0.011 | -0.043 | 0.669 | (-0.059, 0.038) |
| | Recognition | -0.049 | -0.029 | 0.776 | (-0.386, 0.289) |
| Occipital WMH | Delayed Recall | -2.009 | -0.171 | 0.040* | (-0.348, -0.182) |
| | Learning Slope | -0.312 | -0.180 | 0.059 | (-0.636, 0.011) |
| | Percent Forgetting | 0.097 | 0.129 | 0.158 | (-0.038, 0.232) |
| | Retroactive Interference | -0.038 | -0.056 | 0.568 | (-0.169, 0.093) |
| | Recognition | -0.878 | -0.190 | 0.055 | (-1.774, 0.019) |
| Total WMH | Delayed Recall | 0.070 | 0.025 | 0.768 | (-0.396, 0.535) |
| | Learning Slope | 0.039 | 0.096 | 0.324 | (-0.039, 0.117) |
| | Percent Forgetting | -0.002 | -0.012 | 0.897 | (-0.035, 0.030) |
| | Retroactive Interference | -0.003 | 0.016 | 0.825 | (-0.035, 0.028) |
| | Recognition | -0.007 | -0.007 | 0.946 | (-0.223, 0.209) |

Significant values indicated by *.
Regression model: RAVLT measure ~ Regional WMH + age + sex + education + intercept.

**Supplementary Table 2.** Associations between occipital WMH and left/right MTL subregional volumes.

| WMH Distribution | MTL Subregion | *b*-coefficient | Standardized β-coefficient | p-value | 95% CI |
|---|---|---|---|---|---|
| Occipital WMH | Left CA1 | -0.138 | -0.281 | <0.001* | (-0.219, -0.058) |
| | Right CA1 | -0.116 | -0.248 | 0.005* | (-0.196, -0.036) |
| | Left CA2 | 0.001 | 0.057 | 0.560 | (-0.002, 0.004) |
| | Right CA2 | -0.002 | -0.174 | 0.072 | (-0.005, 0.000) |
| | Left CA3 | -0.010 | -0.217 | 0.028* | (-0.019, -0.001) |
| | Right CA3 | $9.8x10^{-5}$ | 0.002 | 0.985 | (-0.010, 0.010) |
| | Left DG | -0.088 | -0.301 | <0.001* | (-0.137, -0.039) |
| | Right DG | -0.075 | -0.263 | 0.003* | (-0.124, -0.025) |
| | Left ERC | -0.041 | -0.199 | 0.032* | (-0.079, -0.004) |
| | Right ERC | -0.055 | -0.222 | 0.021* | (-0.102, -0.009) |
| | Left BA35 | -0.043 | -0.190 | 0.042 | (-0.084, -0.002) |
| | Right BA35 | 0.000 | -0.011 | 0.992 | (-0.39, 0.038) |
| | Left BA36 | -0.250 | -0.338 | <0.001* | (-0.388, -0.112) |
| | Right BA36 | -0.176 | -0.235 | 0.015* | (-0.318, -0.034) |
| | Left SUB | -0.009 | -0.060 | 0.526 | (-0.036, 0.018) |
| | Right SUB | -0.037 | -0.212 | 0.027* | (-0.069, -0.004) |
| | Left PHC | -0.152 | -0.285 | 0.002* | (-0.244, -0.059) |
| | Right PHC | -0.061 | -0.153 | 0.105 | (-0.134, 0.013) |

Significant values indicated by *.
Regression model: Left/Right MTL subregional volume ~ Regional WMH + age + sex + education + intercept.

**Supplementary Table 3.** Associations between MTL subregional volumes and RAVLT measures

| MTL Subregion | RAVLT Measure | *b*-coefficient | Standardized β-coefficient | p-value | 95% CI |
|---|---|---|---|---|---|
| Total CA1 | Delayed Recall | 2.114 | 0.166 | 0.038* | (0.115, 4.113) |
| | Learning Slope | -0.142 | -0.077 | 0.418 | (-0.488, 0.204) |
| | Percent Forgetting | -0.181 | -0.222 | 0.013* | (-0.322, -0.039) |
| | Retroactive Interference | 0.135 | 0.185 | 0.052 | (-0.001, 0.271) |
| | Recognition | 0.690 | 0.151 | 0.126 | (-0.198, 1.578) |
| Total Dentate Gyrus | Delayed Recall | 3.956 | 0.187 | 0.016* | (0.737, 7.176) |
| | Learning Slope | 0.006 | 0.002 | 0.983 | (-0.556, 0.568) |
| | Percent Forgetting | -0.368 | -0.272 | 0.002* | (-0.593, -0.142) |
| | Retroactive Interference | 0.248 | 0.205 | 0.027* | (0.028, 0.468) |
| | Recognition | 1.293 | 0.168 | 0.083 | (-0.170, 2.755) |
| Total Perirhinal Cortex (BA36) | Delayed Recall | 2.445 | 0.285 | <0.001* | (1.263, 3.628) |
| | Learning Slope | 0.011 | 0.009 | 0.918 | (0.203, 0.226) |
| | Percent Forgetting | -0.179 | -0.327 | <0.001* | (-0.263, -0.095) |
| | Retroactive Interference | 0.143 | 0.291 | <0.001* | (0.062, 0.225) |
| | Recognition | 0.653 | 0.214 | 0.018* | (0.115, 1.190) |
| Total Parahippocampal Cortex | Delayed Recall | 2.381 | 0.170 | 0.031* | (0.222, 4.541) |
| | Learning Slope | -0.165 | -0.081 | 0.383 | (-0.540, 0.209) |
| | Percent Forgetting | -0.237 | -0.265 | 0.002* | (-0.387, -0.087) |
| | Retroactive Interference | 0.069 | 0.086 | 0.359 | (-0.080, 0.218) |
| | Recognition | 0.159 | 0.030 | 0.756 | (-0.849, 1.166) |

Significant values indicated by *.

Regression model: RAVLT measure ~ Total MTL subregional volume + age + sex + education + intercept.

**Supplementary Table 4.** Associations between left/right MTL subregional volumes and delayed recall

| MTL Subregion | RAVLT Measure | *b*-coefficient | Standardized β-coefficient | p-value | 95% CI |
|---|---|---|---|---|---|
| Left CA1 | Delayed Recall | 4.112 | 0.170 | 0.036* | (0.270, 7.954) |
| Right CA1 | | 3.808 | 0.151 | 0.056 | (-0.100, 7.717) |
| Left DG | | 7.806 | 0.192 | 0.014* | (1.611, 14.00) |
| Right DG | | 6.750 | 0.166 | 0.032* | (0.575, 12.925) |
| Left BA36 | | 4.048 | 0.253 | <0.001* | (1.837, 6.258) |
| Right BA36 | | 4.496 | 0.279 | <0.001* | (2.252, 6.741) |
| Left PHC | | 4.450 | 0.195 | 0.013* | (0.948, 7.953) |
| Right PHC | | 3.301 | 0.106 | 0.168 | (-1.407, 8.008) |

Significant values indicated by *.
Regression model: Delayed Recall ~ Left/Right MTL subregional volume + age + sex + education + intercept.
